# SARS-CoV-2 mRNA vaccination elicits robust and persistent T follicular helper cell response in humans

**DOI:** 10.1101/2021.09.08.459485

**Authors:** Philip A. Mudd, Anastasia A. Minervina, Mikhail V. Pogorelyy, Jackson S. Turner, Wooseob Kim, Elizaveta Kalaidina, Jan Petersen, Aaron J. Schmitz, Tingting Lei, Alem Haile, Allison M. Kirk, Robert C. Mettelman, Jeremy Chase Crawford, Thi H.O. Nguyen, Louise C. Rowntree, Elisa Rosati, Michael K. Klebert, Teresa Suessen, William D. Middleton, the SJTRC Study Team, Joshua Wolf, Sharlene A. Teefey, Jane A. O’Halloran, Rachel M. Presti, Katherine Kedzierska, Jamie Rossjohn, Paul G. Thomas, Ali H. Ellebedy

**Affiliations:** Department of Emergency Medicine, Washington University School of Medicine, Saint Louis, MO, USA.; Center for Vaccines and Immunity to Microbial Pathogens, Washington University School of Medicine, Saint Louis, MO, USA.; Department of Immunology, St. Jude Children’s Research Hospital, Memphis, TN, USA.; Department of Pathology and Immunology, Washington University School of Medicine, Saint Louis, MO, USA.; Division of Allergy and Immunology, Department of Internal Medicine, Washington University School of Medicine, Saint Louis, MO, USA; Infection and Immunity Program & Department of Biochemistry and Molecular Biology, Biomedicine Discovery Institute, Monash University, Clayton, Victoria 3800, Australia; Clinical Trials Unit, Washington University School of Medicine, Saint Louis, MO, USA.; Department of Microbiology and Immunology, University of Melbourne, at Peter Doherty Institute for Infection and Immunity, Parkville, Victoria, Australia; Institute of Clinical Molecular Biology, Christian-Albrecht University of Kiel, Kiel, Germany; Mallinckrodt Institute of Radiology, Washington University School of Medicine, Saint Louis, MO, USA.; Department of Infectious Diseases, St. Jude Children’s Research Hospital, Memphis, TN, USA.; Division of Infectious Diseases, Department of Internal Medicine, Washington University School of Medicine, Saint Louis, MO, USA.; Institute of Infection and Immunity, Cardiff University, School of Medicine, Heath Park, Cardiff, United Kingdom; The Andrew M. and Jane M. Bursky Center for Human Immunology and Immunotherapy Programs.

**Keywords:** COVID-19, T follicular helper cell, lymph node, CD4^+^ T cell, mRNA vaccination

## Abstract

SARS-CoV-2 mRNA vaccines induce robust anti-spike (S) antibody and CD4+ T cell responses. It is not yet clear whether vaccine-induced follicular helper CD4+ T (TFH) cell responses contribute to this outstanding immunogenicity. Using fine needle aspiration of draining axillary lymph nodes from individuals who received the BNT162b2 mRNA vaccine, we show that frequency of TFH correlates with that of S-binding germinal center B cells. Mining of the responding TFH T cell receptor repertoire revealed a strikingly immunodominant HLADPB1* 04-restricted response to S167-180 in individuals with this allele, which is among the most common HLA alleles in humans. Paired blood and lymph node specimens show that while circulating S-specific TFH cells peak one week after the second immunization, S-specific TFH persist at nearly constant frequencies for at least six months. Collectively, our results underscore the key role that robust TFH cell responses play in establishing long-term immunity by this efficacious human vaccine.

## Introduction

The COVID-19 pandemic necessitated rapid late-stage clinical trials of mRNA vaccine technology (Anderson et al., 2020; Baden et al., 2021; Jackson et al., 2020; Polack et al., 2020; Verbeke et al., 2021; Walsh et al., 2020; Widge et al., 2021) that resulted in the first FDA-approved vaccine using this technology platform. The two mRNA vaccines developed by Pfizer/BioNTech (BNT162b2) (Polack et al., 2020) and Moderna (mRNA-1273) (Baden et al., 2021) have proven instrumental in the initiation of widespread vaccination campaigns in the United States and around the world. Both vaccines engender very high-titer circulating anti-SARS-CoV-2 Spike (S) protein-specific antibodies that can neutralize the originally circulating SARS-CoV-2 strain (Jackson et al., 2020; Walsh et al., 2020) as well as other variants that have emerged since the vaccine design phase (Chen et al., 2021; Wang et al., 2021a, 2021b; Wu et al., 2021). Neutralizing antibodies induced by mRNA vaccines appear to be the key correlate of protection from COVID-19 in animal models (Corbett et al., 2021) and in humans (Khoury et al., 2021). The COVID-19 mRNA vaccines exhibit the highest efficacy in phase 3 studies among widely utilized COVID-19 vaccines worldwide (Al Kaabi et al., 2021; Baden et al., 2021; Logunov et al., 2021; Polack et al., 2020; Sadoff et al., 2021; Voysey et al., 2021). Understanding exactly how mRNA vaccines elicit such robust and protective immune responses in humans is necessary for the extension and application of this technology to novel vaccines against other important pathogens.

The lymph node germinal center (GC) reaction is critical to the development of long-lasting, high-affinity antibody responses (Ripperger and Bhattacharya, 2021; Victora and Nussenzweig, 2012). T follicular helper (TFH) cell responses are necessary for forming and sustaining GC reactions and for the development of both long-lived plasma cells and memory B cells (Crotty, 2011; Qi, 2016; Ueno et al., 2015). Detailed analysis of vaccination induced GC reactions in humans is just beginning to be explored with recently published studies that have sampled the draining axillary lymph node using serial fine needle aspiration (FNA) following intramuscular deltoid vaccination (Turner et al., 2020, 2021). Importantly, it appears that the GC reaction in humans persists over a longer period of time (Turner et al., 2020, 2021) than what was anticipated from studies in mice (Good-Jacobson et al., 2014; Weisel et al., 2016). Determining the epitope targets and dynamics of SARS-CoV-2-specific TFH cells induced in human draining lymph nodes during an active immune response is critical for understanding the role of TFH in the development of long-lived plasma cells and memory B cells following vaccination.

## Results

### Human TFH population size mirrors the size of the antigen-specific GC B cell population

We conducted a prospective observational study to follow vaccine-induced immune responses in a cohort of 41 healthy adults who received the BNT162b2 mRNA vaccine (Turner et al., 2021). Demographics of the full cohort have been previously reported (Turner et al., 2021). Fourteen members of the cohort underwent axillary lymph node FNA. All subjects were vaccinated with two 30 µg doses of BNT162b2 approximately twenty-one days apart. Blood and/or FNA samples were obtained at day 0 (prior to the first vaccine dose), day 21 (immediately prior to the second vaccine dose), day 28, day 35, day 60, day 110 and day 200 according to the schedule listed in Figure 1A. This manuscript reports exclusively on the 14 subjects who underwent lymph node FNA. Demographics of the included individuals are listed in Table 1. None of the included subjects reported previous infection with SARS-CoV-2.

**Figure 1.**
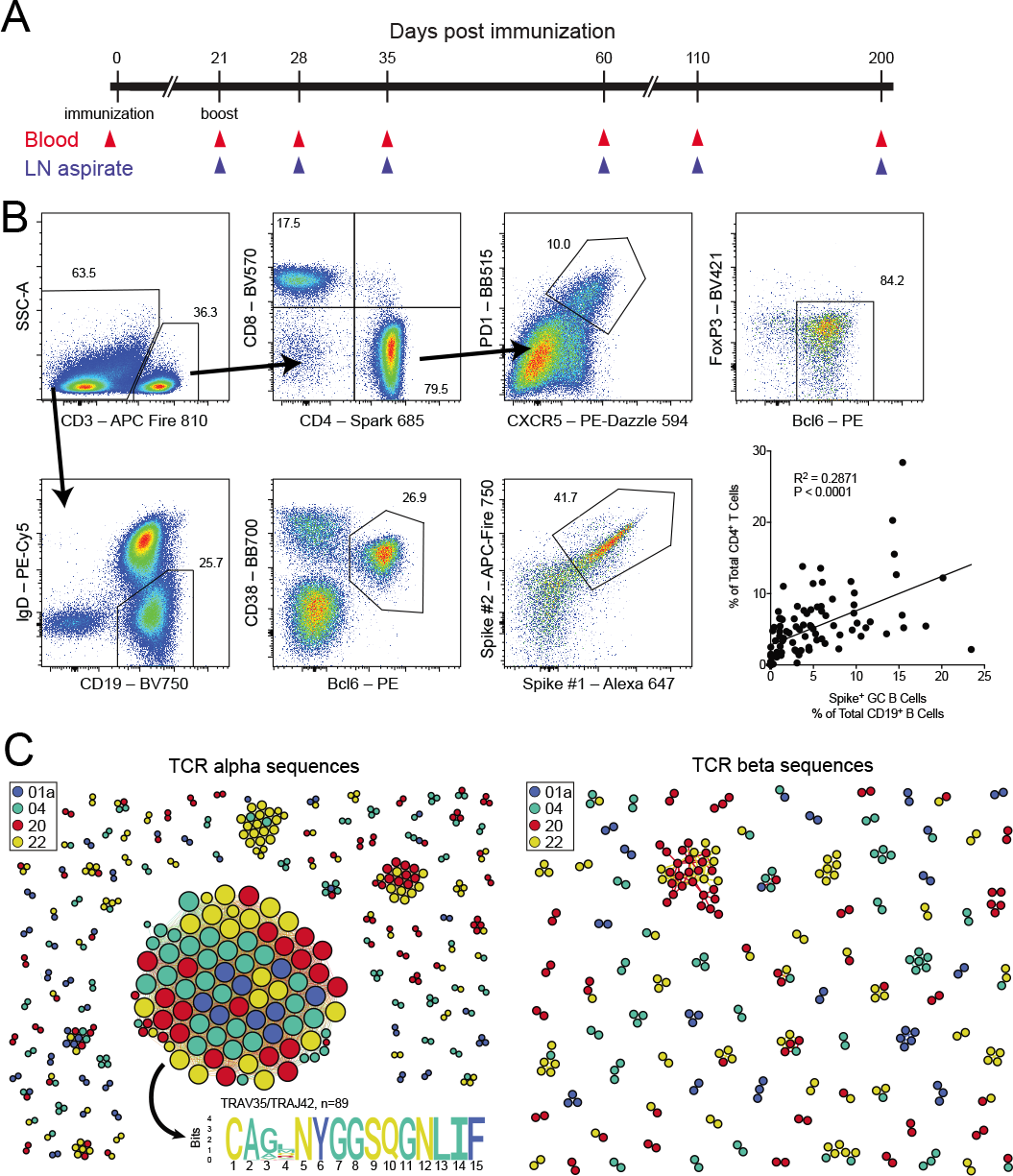
Human lymph node TFH response following mRNA vaccination. (**A**) Study timeline. Day 0 blood samples were obtained prior to the first dose of the vaccine and day 21 samples were taken prior to the second dose of the vaccine. (**B**) Total lymph node TFH (CD3^+^CD4^+^CXCR5^+^PD1^+^Bcl-6^+^FoxP3^-^) measured as the frequency of total lymph node CD4^+^ T cells were compared to the total frequency of lymph node germinal center B cells (CD19^+^IgD^low^Bcl-6^+^CD38^int^) that positively stained with two separate fluorescently-labeled Spike protein probes using linear regression. N=96 individual lymph node samples obtained from all 14 study subjects between and including study days 21 and 200. (**C**) Similarity network of the 500 most abundant TCR α sequences (left) and TCR β sequences (right) from the lymph node TCR repertoire obtained from sorted TFH of 4 individual donors (01a, 04, 20 and 22) 60 days after mRNA vaccination. Each vertex corresponds to an individual TCR clonotype, which are connected to adjacent data points if they have identical VJ-segments and less than 2 mismatches in the CDR3 amino acid sequence. The size of the vertex corresponds to the vertex degree (number of neighbours).

**Table 1.**

| **Study ID** | **Sex** | **Race** | **Ethnicity** | **Age** | **Number of**<br>**Lymph**<br>**Nodes**<br>**Sampled** |
| --- | --- | --- | --- | --- | --- |
| 01a | Male | White | Non-hispanic | 34 | 1 |
| 02a | Male | White | Hispanic | 37 | 2 |
| 04 | Female | White | Non-hispanic | 38 | 2 |
| 07 | Female | White | Non-hispanic | 33 | 1 |
| 08 | Female | White | Non-hispanic | 27 | 1 |
| 10 | Female | White | Non-hispanic | 27 | 2 |
| 13 | Male | White | Non-hispanic | 34 | 1 |
| 15 | Female | Black | Non-hispanic | 52 | 2 |
| 16 | Male | White | Non-hispanic | 37 | 2 |
| 20 | Female | White | Non-hispanic | 48 | 2 |
| 21 | Female | White | Non-hispanic | 31 | 1 |
| 22 | Male | White | Non-hispanic | 36 | 1 |
| 28 | Female | Asian | Non-hispanic | 44 | 1 |
| 43 | Male | Asian | Non-hispanic | 40 | 1 |

To estimate the role of TFH cells in the generation of B cells specific for the SARS-CoV-2 S protein, we analyzed the frequency of the S-specific GC B cell response (defined as S-specific CD19^+^IgD^low^Bcl-6^+^CD38^int^ B cells) among all lymph node resident B cells and the frequency of total lymph node-resident CD4^+^ T cells that exhibited a TFH cell phenotype (Bcl-6^+^CXCR5^+^PD1^+^FoxP3^-^) in lymph node samples taken from each of the 14 individuals. We evaluated all FNA samples obtained between 21 and 200 days following primary vaccination. Six of the fourteen subjects underwent repeated sampling of two separate axillary lymph nodes. We found a strong positive correlation between the size of the S-specific GC B cell population in the lymph node and the total TFH cell population frequency (Figure 1B).

### Discovery and characterization of an immunodominant DPB1*04:01-restricted CD4^+^ T cell population

We next sought to illuminate the antigen-specificity of the characterized TFH population that is so closely correlated with the size of the S-specific GC B cell population. To do this, we sorted total TFH from FNA samples obtained on day 60 from four separate subjects and reconstructed their T cell receptor (TCR) repertoires using unpaired sequencing of the TCR α and TCR β chains. Surprisingly, clonally expanded TCRs formed a prominent α chain cluster that was shared among all 4 donors (Figure 1C). We did not observe a similar shared cluster in the TCR β chain repertoires (Figure 1C). We observed the same α motif in a previously published paper (Minervina et al., 2021), where it was the largest signal and corresponded to 0.2% of total CD4^+^ T cells and 16.3% of estimated SARS-CoV-2-responding CD4^+^ T cells in the blood at the peak of the acute response. Large clusters of TCRs with sequence similarity is an indication of convergent selection of similar receptors to the same antigen (Dash et al., 2017; Glanville et al., 2017; Pogorelyy et al., 2019). As this motif was present among expanded clones in many donors, it likely recognizes an immunodominant epitope from SARS-CoV-2 presented in the context of a common HLA class II allele.

In order to decode the specificity of the heterodimer αβTCR, we first needed to determine what β chains pair with the TCRα chain motif that we identified (Figure 2A). To do this, we queried publicly available CD4^+^ paired TCR datasets. We used two datasets that have paired αβTCR sequences from CD4^+^ T cells after antigen-reactive T cell enrichment following stimulation with SARS-CoV-2 peptides (Bacher et al., 2020; Meckiff et al., 2020). We searched for our CDR3α motif (“CA[G/A/V]XNYGGSQGNLIF”) in these datasets and found 1329 out of 44256 unique TCRs in Bacher et al, but only 53 out of 43745 in Meckiff et al with the matched CDR3α motif. We next used the identified β chains to look for overlap in the MIRA dataset (Nolan et al., 2020) – a large dataset produced by Adaptive Biotech linking TCR sequences to SARS-CoV-2 epitopes. We identified 64 TCRs from Bacher et al highly similar (up to one amino acid mismatch in CDR3, identical CDR1 and CDR2) to MIRA TCRs reactive to the overlapping peptide pool from SARS-CoV-2 Spike protein 160-218 positions (Figure 2B). Interestingly, this part of the spike protein was not used for stimulation in Meckiff et al, explaining why we found only a small number of TCRs of interest in this dataset and indirectly supporting the predicted identification of the peptide from the MIRA dataset.

**Figure 2.**
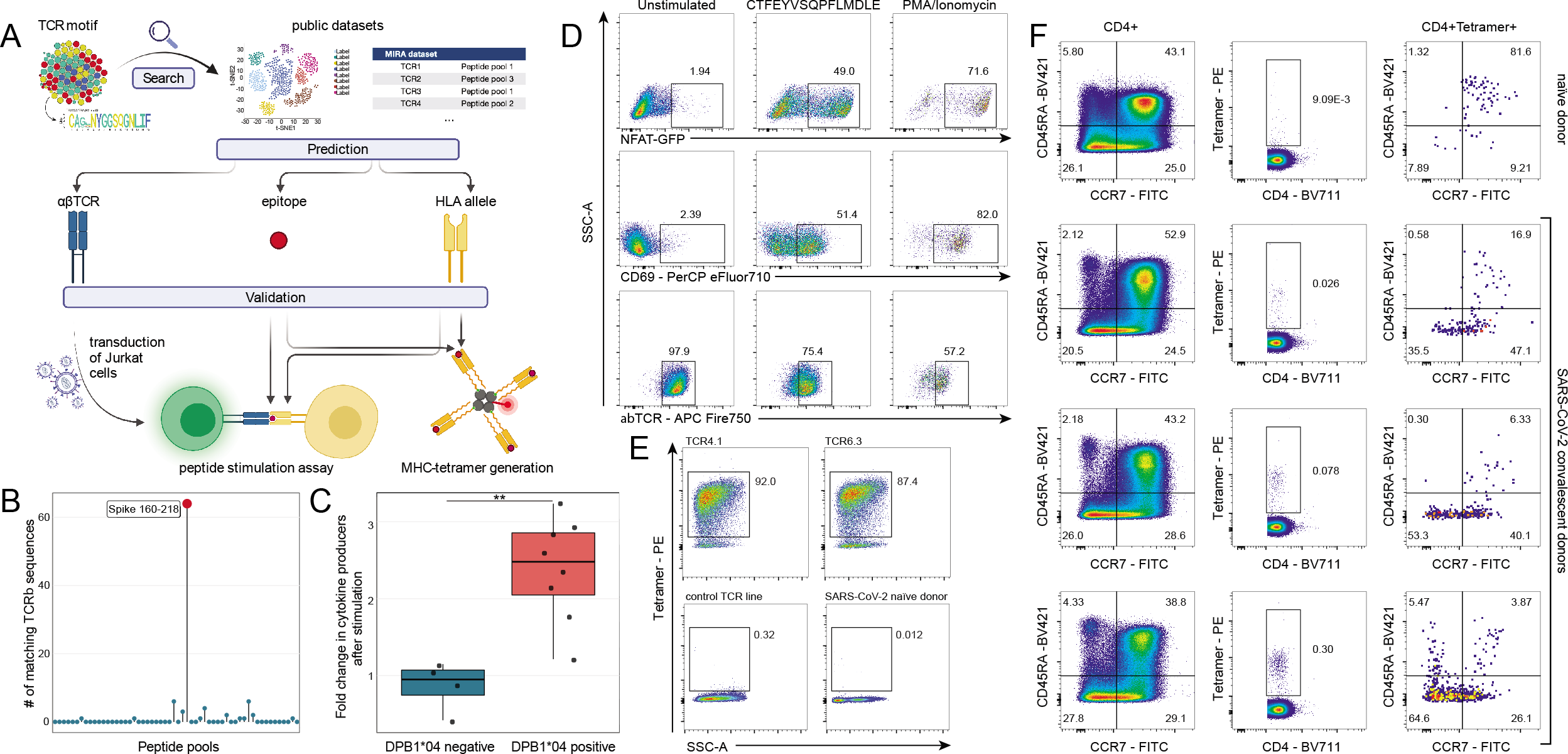
S167-180 epitope discovery and HLA class II tetramer validation. **(A)** Response identification process. The identified TCRα motif of interest was used to query large public scRNAseq datasets (Bacher et al., 2020; Meckiff et al., 2020) to identify potential partner TCRβ chains and then matched to the large MIRA dataset that used TCRβ sequencing (Nolan et al., 2020) to predict HLA-restriction and cognate epitope. To validate our prediction, we generated a T cell line expressing the putative αβTCR and we also generated HLA class II tetramers. **(B)** Identification of peptide pool for the motif TCRs using the MIRA dataset. TCRβ chains from paired αβTCRs with CDR3α motif (CA[G/A/V].NYGGSQLIF) were searched in the MIRA dataset allowing for up to one mismatch in CDR3 amino acid sequence. The Y-axis shows the number of TCRβ chains from Bacher et al. matching to TCRβ from different MIRA SARS-CoV-2 peptide pools. Largest hit (red dot) corresponded to the peptide pool spanning amino acid positions 160-218 from S-protein. **(C)** Average fold change in CD4^+^/CD69^+^ T cells (producing IL2, TNFα, or IFNγ) per 10^6^ cells following CTFEYVSQPFLMDLE peptide stimulation of DPB1*04-positive and -negative SJTRC PBMCs. PBMCs collected during SARS-CoV-2 convalescence or post-vaccination with BNT162b2 were used for intracellular cytokine staining assay. Average fold changes were compared using a Mann-Whitney *U* test; p =0.004. Gating strategy is shown in Supplementary Figure 1. (D) Jurkat cell line expressing the predicted TCR after stimulation with the predicted epitope. Left column: negative control; middle column: TCR4.1 cell line co-cultured with PBMCs from healthy DPB1*04:01-positive donor pulsed with CTFEYVSQPFLMDLE peptide (S167-180); right column: positive control. Top row: NFAT-GFP reporter expression. Middle row: CD69 surface expression. Bottom row: Downregulation of the TCR on cell surface. **(E)** S167-180 tetramer staining identifies epitope specific T cells with high specificity. Top row: staining of TCR4.1 and TCR6.3 Jurkat cell lines. Bottom left, staining of Jurkat cell line expressing TCR with other known specificity, bottom right: staining of PBMCs from SARS-CoV-2 naive individual. **(F)** S167-180 tetramer^+^ cells have predominantly effector memory phenotype in COVID-19 convalescent patients. Each row represents an individual donor. Left column: CCR7 and CD45RA distribution in bulk CD3^+^CD4^+^ cells. Middle column: S167-180 tetramer staining of CD3^+^CD4^+^ cells. Right column: memory/naive phenotypes of CD3^+^CD4^+^S167-180 tetramer^+^ cells. Gating strategies for **(D)**, **(E)**, **(F)** are shown in Supplementary Figure 2.

Five of six subjects recognizing this peptide pool in the MIRA database had available HLA-typing. These five shared the DPB1*04:(01/02) and DQB1*06:(02/03) alleles. To establish HLA-restriction of the response of interest and to narrow the search to a single peptide, we next used NetMHCII2.3 (Jensen et al., 2018) to look for predicted epitopes from the S160-218 peptide pool that are presented by one or both of these shared alleles. We found that peptides containing the core sequence **YVSQPFLMD** were predicted to strongly bind the DPB1:04:01 and DPB1:04:02 alleles, while no strong binders were identified for the DQB1*06:(02/03) alleles. Interestingly, TCR epitopes with this core sequence (**YVSQPFLMD**, S170-178) have been previously described in prominent epitope discovery studies (Peng et al., 2020; Tarke et al., 2021), where the response was identified in multiple donors. However, this response has not previously been reported to be HLA-DPB1*04-restricted. As an initial investigation of this possible HLA-restriction, we obtained post-vaccination peripheral blood from participants in the ongoing SJTRC study (SJTRC, NCT04362995). PBMCs from these participants were stimulated with purified S166-180 peptide (CTFE**YVSQPFLMD**LE) and the responses were measured by intracellular cytokine staining and flow cytometry. We determined that participants with the HLA-DPB1*04 allele had increased total cell counts of CD4^+^CD69^+^ T cells producing IL-2, TNFα, or IFNγ compared to participants without this allele (Figure 2C, Supplementary Figure 1). We then moved forward with more rigorous experimental validation of our paired TCR, peptide epitope, and restricting HLA combination (Figure 2A).To do this, we selected two paired TCRs from Bacher et al that included the same TCRα, but different TCRβ chains that we designated TCR4.1 and TCR6.3. We transduced these each into separate Jurkat TCR-negative cell lines that also express an endogenous NFAT-GFP reporter to allow for tracking intracellular signaling downstream of the transduced paired TCR following TCR engagement. We demonstrated epitope recognition of the CTFE**YVSQPFLMD**LE peptide through the stimulated transduced TCR in assays using these Jurkat cell lines co-cultured with PBMC from an HLA-DPB1*04^+^ donor as APCs (Figure 2D, Supplementary Figure 2).

We next generated HLA class II tetramer to probe the antigen-specific T cell response we discovered. We tested our HLA-DPB1*04 S167-180 tetramer using the two transduced TCR4.1 and TCR6.3 Jurkat cell lines and showed high sensitivity and low-background staining (Figure 2E). We then used the S167-180 tetramer to look for antigen-specific CD4^+^ T cells in PBMC from three HLA-DPB1*04^+^ SARS-CoV-2 convalescent donors and in a control HLA-DPB1*04^+^ SARS-CoV-2 naïve donor. We found a small number of tetramer-specific cells predominantly in the naïve subpopulation (CCR7^+^CD45RA^+^) in the naïve donor, and a much larger number of tetramer-specific cells that were primarily effector memory (CCR7^-^CD45RA^-^) in the SARS-CoV-2 convalescent donors (Figure 2F). We then sequenced tetramer-specific TCRs from these convalescent donors using our previously described scTCRseq approach (Wang et al., 2012). The majority (64%) of sequenced cells had the same TRAV35-CA[G/A/V]NYGGSQGNLIF TCRα motif that we initially identified, and >80% of all sequences included TRAV35, suggesting that the discovered TCRα motif is the most frequent mode of recognition for this epitope (**Supplementary Table 1**).

### Tracking S167-180 antigen-specific CD4^+^ T cell responses in blood and draining lymph nodes following BNT162b2 vaccination

With the discovery of an immunodominant SARS-CoV-2-S epitope restricted by the HLA-DPB1*04:01 allele that is found at high frequency (>40%) in many populations around the world (allelefrequencies.net), we used the S167-180 HLA class II tetramer to evaluate the 14 individuals with available lymph node FNA samples to empirically determine which individuals were HLA-DPB1*04:01^+^ and thus made the S167-180-specific CD4^+^ T cell response. Nine of 14 FNA subjects made a detectable S167-180-specific response in peripheral blood following boost vaccination. We next tracked and characterized this response over time in frozen PBMC (N=7 subjects) and frozen lymph node FNA samples (N=3 subjects) from a convenience sample of the 9 subjects with sufficient samples remaining for analysis. The S167-180-specific CD4^+^ T cell response peaked in peripheral blood 28 days after primary vaccination, 7 days after vaccine boost, and remained present in the blood at detectable frequencies through the entire study interval (Figure 3A and 3B). Most S167-180-specific CD4^+^ T cells circulating in peripheral blood exhibited a CD45RO^+^CCR7^-^ effector memory surface phenotype similar to what we observed in SARS-CoV-2 convalescent donors (Figure 3C). A subset of tetramer-positive CD4^+^ T cells in the first 35 days following primary vaccination exhibited an activated surface phenotype characterized by upregulation of both CD38 and HLA-DR (Figure 3D). This activated CD4^+^ T cell phenotype disappeared by day 60 post-primary vaccination. Most circulating S167-180-specific CD4^+^ T cells expressed both PD1 and ICOS at high levels on days 21 and 28 following primary vaccination with a gradual decrease in the mean fluorescent intensity of PD1 and ICOS throughout the remaining study interval to a level more consistent with that found on the majority of circulating CD4^+^ T cells in line with resolution of T cell activation (Figure 3E). A subset of S167-180-specific CD4^+^ T cells accounting for approximately 5-15% of the total number of circulating S167-180-specific CD4^+^ T cells exhibited the CXCR5^+^PD1^+^ circulating TFH phenotype (Figure 3F). These circulating S167-180-specific TFH cells peaked 28 days after primary vaccination, 7 days after vaccine boost and then decreased over time, becoming difficult to detect in the blood of some subjects by the final study time-point (Figure 3G).

**Figure 3.**
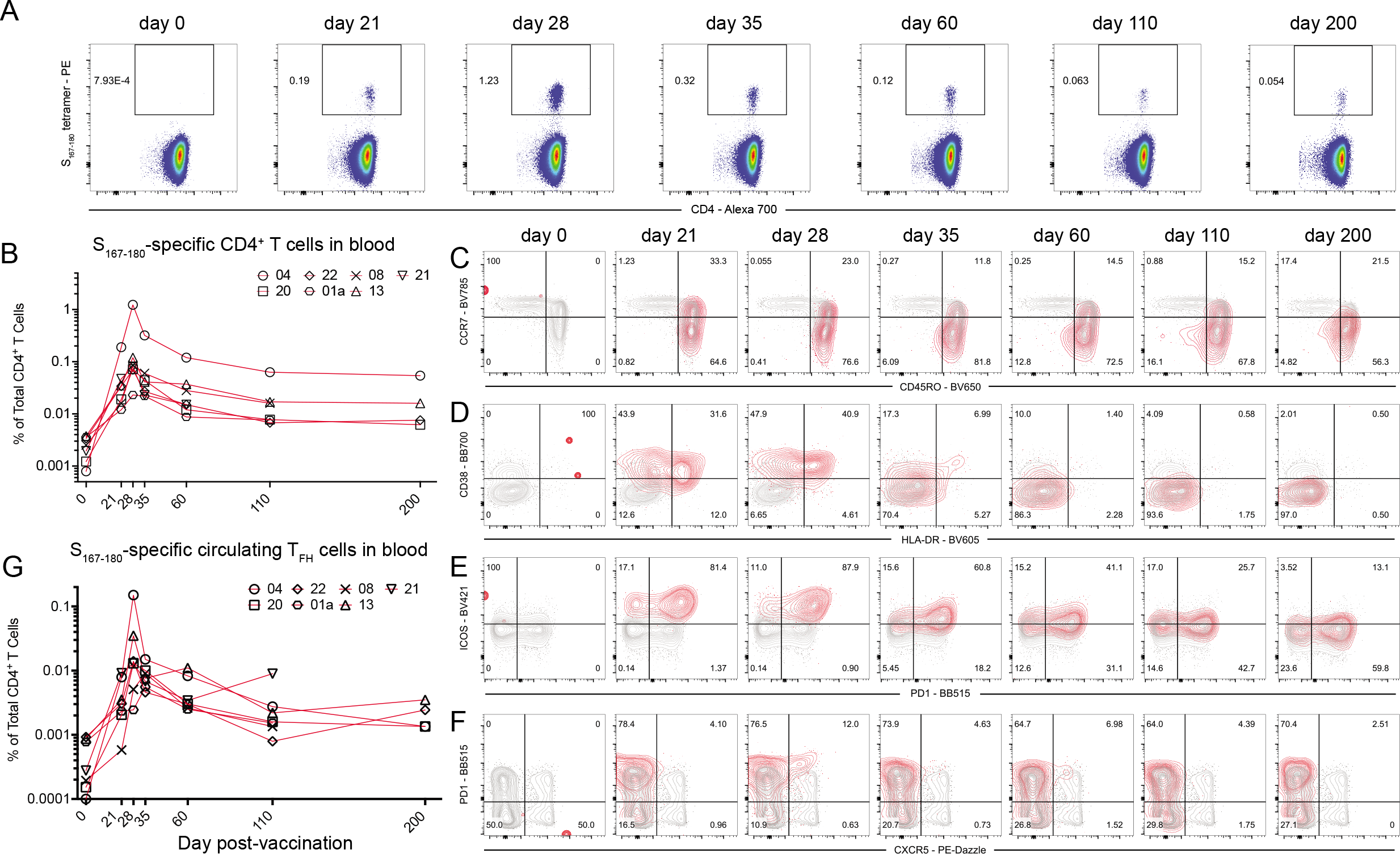
S167-180 response in peripheral blood following BNT162b2 vaccination. (**A**) Representative flow cytometry plots of S167-180 tetramer staining following vaccination of subject 04. Frequency displayed is the percent of live CD3^+^CD4^+^ T cells in the blood that are tetramer positive. (**B**) The frequency of S167-180 tetramer^+^ cells in the blood over time in 7 of the study subjects with available PBMC from most time-points. (**C-F**) Surface phenotype of circulating S167-180 tetramer^+^ cells over time. Representative flow cytometry overlay plots from subject 04 showing total CD4^+^ T cell (grey contours) and tetramer positive (red contours) populations. (**C**) The majority of S167-180 tetramer^+^ cells retain an “effector memory” (CD45RO^+^CCR7^-^) surface phenotype following vaccination. (**D**) A subset of S167-180 tetramer^+^ cells undertake an “activated” surface phenotype (HLA-DR^+^CD38^+^) in the two weeks following vaccination. (**E**) ICOS and PD-1 are upregulated on the majority of S167-180 tetramer^+^ cells prior to and 7 days following boost vaccination. (**F**) A small subset of S167-180 tetramer^+^ cells undertake a “circulating TFH” surface phenotype (CXCR5^+^PD1^+^) following boost vaccination, but the majority of circulating S167-180 tetramer^+^ cells do not exhibit this phenotype. (**G**) S167-180 tetramer^+^CXCR5^+^PD1^+^ cells as a percentage of total live CD3^+^CD4^+^ T cells over time.

In contrast to circulating TFH cells, the frequency of S167-180-specific CD4^+^ T cells remained high in the draining axillary lymph node through at least day 60 following primary vaccination and persisted at a relatively constant and high frequency in one of the three study subjects through day 200 following primary vaccination – more than 170 days following vaccine boost (Figure 4). The prolonged persistence of S-specific TFH that we report here in the draining axillary lymph nodes corresponds well with the long-lived germinal center B cell responses recently reported in the same cohort of subjects (Turner et al., 2021). The vast majority of S167-180-specific CD4^+^ T cells in lymph node FNA samples co-expressed CXCR5 and PD1, surface markers of TFH cells (Figure 4A). Furthermore, the frequency of S167-180-specific CD4^+^ T cells in the FNA samples remained constantly high or even increased as the frequency of S167-180-specific CD4^+^ T cells in the peripheral blood contracted (Figure 4B).

**Figure 4.**
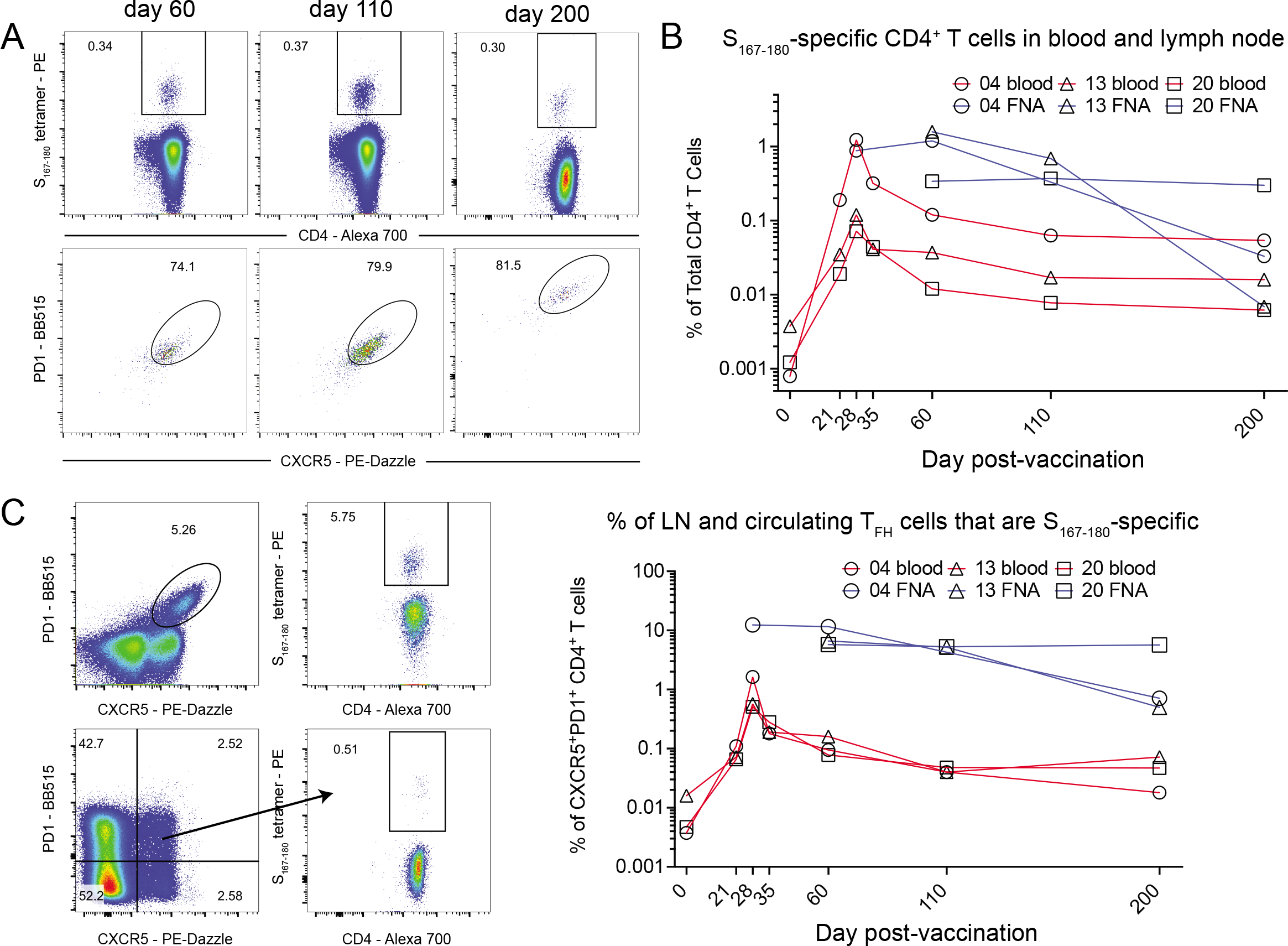
S167-180 response in the draining lymph node following BNT162b2 vaccination. (**A**) Representative flow cytometry plots of subject 20 demonstrating the frequency of S167-180 tetramer^+^ cells expressed as a percentage of total CD4^+^ T cells in the lymph node FNA sample (top row). The bottom row demonstrates CXCR5 and PD1 surface expression on the gated S167-180 tetramer^+^ cells from the row above. (**B**) The percentage of total CD4^+^ T cells that are S167-180 tetramer^+^ in blood (red lines) and FNA (blue lines) in matched samples taken at the same time-points from subjects with available sample (circle – 04, triangle – 13, square -20). (**C**) The percentage of CXCR5^+^PD1^+^ T cells that are S167-180 tetramer^+^ over time in both the blood (red lines) and FNA (blue lines).

We next examined the frequency of S167-180-specific CD4^+^ T cells in the total CXCR5^+^PD1^+^ TFH population in both the blood and the lymph nodes over time. We found that this population rapidly expanded in the blood – peaking at day 28 after primary vaccination, 7 days after vaccine boost – and then became challenging to detect by days 110 and 200 (Figure 4C) as we had previously noted when examining this population as a proportion of total CD4^+^ T cells in Figure 3G. In contrast, the frequency of the S167-180-specific TFH population remained nearly constant within the total TFH population over time in the lymph node – until this response also disappeared and was difficult to detect at day 200 in 2 of the 3 subjects with FNA sample available for analysis (Figure 4C). Together, these results demonstrate that a small subset of clonally-identical antigen-specific CD4^+^ T cells circulating in peripheral blood following vaccination develop a surface phenotype consistent with circulating TFH cells. This coincides with the development of clonally-identical TFH cells in the draining lymph node. Furthermore, while this population nearly disappears from circulating blood 110 days after vaccination, the response remains nearly constant in the lymph node in the presence of an ongoing germinal center reaction. Overall, our findings are consistent with the development of diverse lineages of effector CD4^+^ T cells – those that express a surface phenotype consistent with TFH and those that do not - from a single population of clonally-identical naïve CD4^+^ T cells. This is consistent with observations in mouse models where the duration of the T cell receptor / peptide / MHC class II interaction correlated with the overall balance between Th1 and TFH cell frequency (Tubo et al., 2013).

### Diverse clonal populations of TFH in the human GC persist at a consistent frequency over time

We next quantified the contribution of the S167-180 TFH population to the broader clonotypic diversity of the TFH population found in the lymph node from four of the subjects by further analyzing the TCR sequencing data from sorted TFH cells generated for Figure 1C. The clonotypes that compose the S167-180 response made up the largest percentage of total clonotypes present in the lymph node for three of the four subjects and composed the second highest percentage of clonotypes in the fourth subject (Figure 5A). This underscores the importance of the immunodominant HLA-DPB1*04-restricted S167-180 response in the total SARS-CoV-2-specific TFH response of HLA-DPB1*04^+^ vaccinees, who make up approximately 40-50% of the world’s population.

**Figure 5.**
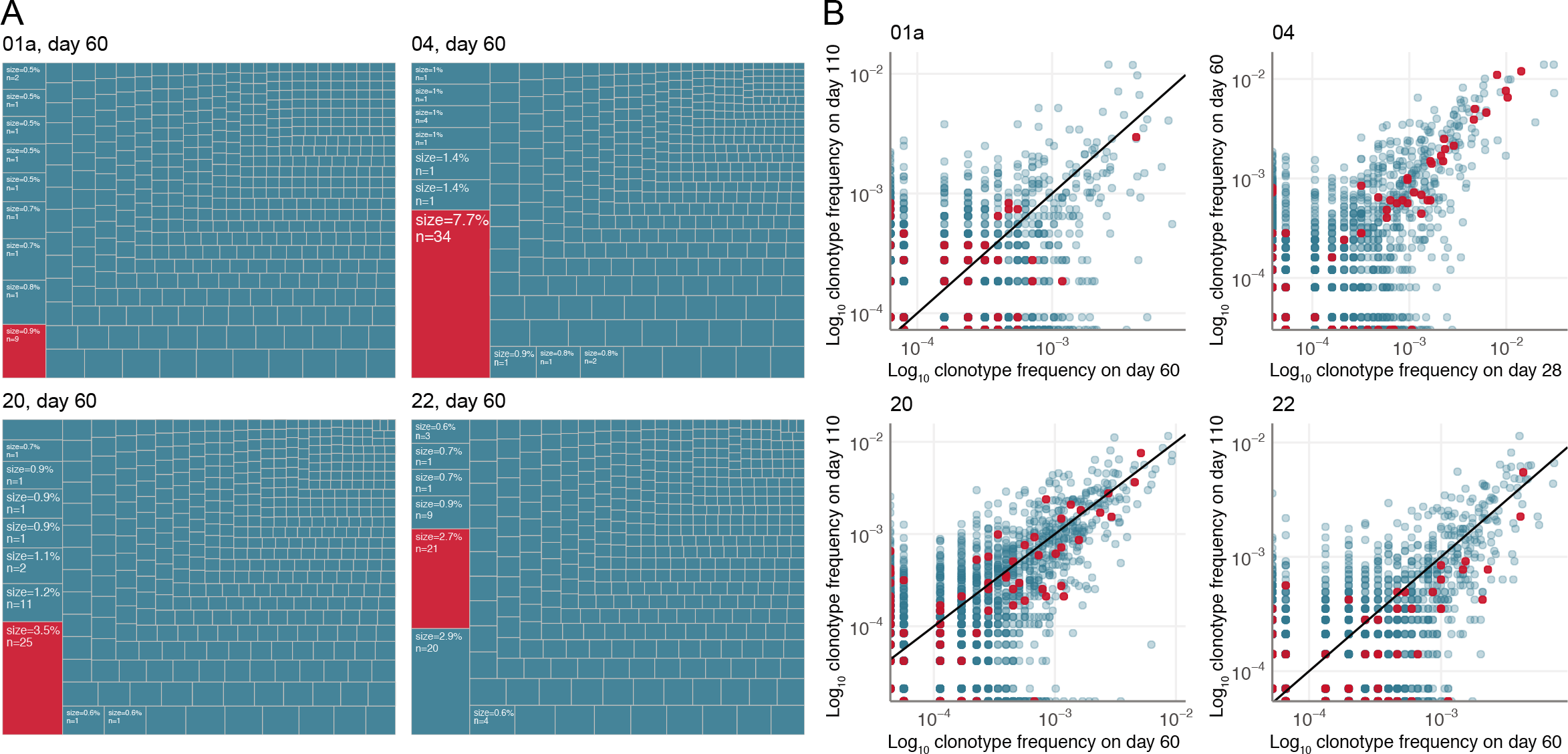
The S167-180 response composes a large fraction of the TFH repertoire that maintains a consistent frequency over time. **(A)** Abundance of the S167-180-specific clones (red boxes) in the lymph node TFH repertoires of 4 donors on day 60 after mRNA vaccination. Listed frequency is the frequency of the examined clonal group (defined as a cluster from Figure 1C) out of total clonal sequences in the sorted TFH sample. The S167-180 response is the largest TFH response in 3 of the 4 examined HLA-DPB1*04:01^+^ subjects lymph nodes. **(B)** Clonotype frequencies of sequenced sorted CXCR5^+^PD1^+^ TFH repertoires from lymph nodes sampled at two separate time points. Each dot corresponds to an individual TCRα clonotype. Frequencies are shown in log scale. Red dots correspond to S167-180-specific clones based on the known alpha chain motif.

To elucidate the clonal composition of the TFH response over time, we sequenced samples from two time-points that were available from these individuals. Three subjects were sequenced at day 60 and day 110 post primary vaccination and one subject was sequenced at day 28 and day 60 following primary vaccination. Three of the subjects exhibited evidence of ongoing antigen-specific TFH responses associated with germinal center responses at all tested time-points by flow cytometry (Figure 4), while one of the four subjects did not have sufficient remaining samples to be analyzed by flow cytometry. Supporting our observations in the flow cytometry analysis of the S167-180 population, we found a strong positive correlation between the frequency of a large number of the TCR α clonotype sequences at the two time points (Figure 5B), including the known S167-180-specific TCR clonotypes (Figure 5B, red data points). This was especially true of the clonotypes found at the highest frequency in each FNA sample which are those that are most likely to represent antigen-specific clonotypes due to their increased presence in the lymph node following vaccination. This positive correlation means that many of these clonotypes were found at the same frequency at both tested time-points. Therefore, the nearly constant frequency of the antigen-specific TFH response over time that we observed in the S167-180-specific response (Figure 4C) is generalizable to other clonally-related and presumably antigen-specific TFH populations in the human lymph node following BNT162b2 vaccination. Our data support a model whereby the human germinal center TFH response is essentially an “on or off”-response in the setting of an active and ongoing germinal center reaction, rather than a response that peaks or dynamically changes in frequency over time.

## Discussion

In this report, we show that the BNT162b2 COVID-19 mRNA vaccine induces robust and persistent TFH responses in the draining lymph nodes of vaccinated individuals. Indirect evidence has existed for some time that robust CD4^+^ T cell responses are required for the generation of high-titer neutralizing antibody responses following COVID-19 infection or mRNA vaccination. This includes data showing a lack of seroconversion in individuals with uncontrolled HIV and extremely low CD4^+^ T cell counts during vaccination (Touizer et al., 2021) as well as several reports that have demonstrated a lack of seroconversion to the standard two-dose BNT162b2 regimen in individuals subjected to T cell-focused immunosuppressive regimens following solid organ transplantation (Kamar et al., 2021). Our current results provide strong and direct evidence that a high-magnitude, antigen-specific CD4^+^ T cell response in the draining lymph nodes is present during the development of high-titer neutralizing antibody responses in the setting of COVID-19 mRNA vaccination.

The temporal relationship we observe between the early appearance and then disappearance of S167-180-specific CD4^+^ T cells exhibiting a circulating TFH phenotype in the blood at the same time that we observe TFH cells in the draining lymph node suggests a complex relationship between these two populations of cells. Our present data support a model of human TFH cell development whereby phenotypically heterogeneous, or even plastic, antigen-specific CD4^+^ T cell populations induced by primary vaccination are activated and expand in the lymph node and circulating compartments prior to the development and migration of more specialized subpopulations that co-express CXCR5 and PD1 to the lymph node GC (Crotty, 2018). In our S167-180 tetramer data, most S167-180-specific CD4^+^ T cells in the blood do not exhibit a circulating TFH phenotype even at the day 28 post-primary vaccination peak of circulating S167-180-specific TFH. Very few, if any, maintained S167-180-specific memory CD4^+^ T cells in the blood express both CXCR5 and PD1. Nevertheless, S167-180-specific TFH cells compose the largest or second largest S-specific TFH population in the lymph node of all evaluated subjects despite the absence of these cells in the circulating blood at the same late time-points. Together, our data support a model whereby clonal populations of circulating CD4^+^ T cells develop into many different lineages, including the TFH cell lineage. Furthermore, we were unable to find a strong direct relationship between the cells known as circulating TFH (circulating antigen-specific CD4^+^CXCR5^+^PD1^+^ cells) and the presence of large populations of clonally-matched antigen-specific TFH participating in an ongoing GC in the lymph node. This is in contrast to data from a study of matched tonsil and blood samples in subjects who were not recently vaccinated or infected where they found substantial clonal overlap between tonsil TFH populations and circulating TFH populations but little overlap between tonsil TFH populations and circulating non- TFH populations (Brenna et al., 2020). Further studies are required to determine the relationship between populations of circulating and lymph node resident TFH cells in both the steady state and following vaccination as these systems are quite distinct.

The discovered DPB1*04 S167-180 response is notable for the extraordinary restriction of the TCRα sequence. This single motif is immediately obvious with even cursory inspection of bulk CD4^+^ TCR sequences from vaccinated or infected individuals. Surprisingly, no prominent TCRβ motif is observed in any of our sequencing of this response, emphasizing the importance of the alpha chain in certain instances of specific epitope recognition (Dash et al., 2017; Minervina et al., 2020; Shomuradova et al., 2020). The high prevalence of DPB1*04 in worldwide populations means that this response is likely immunodominant across multiple populations and contributes significantly to the measured responses in many studies, though its restriction has not been previously assigned. Thus far, none of the prevalent variant SARS-CoV-2 strains has acquired stable mutations in this peptide sequence.

Our study has several limitations, including the small number of subjects and the lack of comprehensive epitope mapping beyond the immunodominant response we identified. There are several questions that we did not address that will be useful topics for future studies, including the extent of clonal overlap between the blood and lymph node CD4^+^ T cell compartments, and the transcriptional profiles of the lymph node TFH response over the long period of clonal stability.

In conclusion, we find that mRNA vaccine technology has an exceptional ability to induce high-frequency antigen-specific B cell (Turner et al., 2021) and antigen-specific CD4^+^ TFH cell responses in the human lymph node following prime-boost administration. These characteristics underlie the development of high titer neutralizing antibodies and protection from infection in vaccinated individuals. The selective enhancement of lymph node TFH responses induced by vaccine regimens represents a broad strategy for improving future vaccines.

## Acknowledgments

The authors would like to thank the study participants for their invaluable contribution to this work. We also thank Greig Lennon from St. Jude Immunology flow core for his help with FACS, and Hartwell Center at St. Jude Children’s Research Hospital for high-throughput sequencing, and Carmen Llerna for assistance in HLA-DP4 protein purification. This study was funded in part by the Washington University Institute for Clinical and Translational Sciences grant UL1TR002345 from the National Center for Advancing Translational Sciences (NCATS) of the National Institutes of Health (NIH) (PAM). JR is supported by an Australian Research Council Laureate Fellowship. KK was supported by the NHMRC Leadership Investigator Fellowship (#1173871) and THON was supported by the NHMRC Emerging Leadership Level 1 Investigator Fellowship (#1194036). This work was funded by ALSAC at St. Jude, the Center for Influenza Vaccine Research for High-Risk Populations (CIVR-HRP) contract number 75N93019C00052 (PGT and KK), the St. Jude Center of Excellence for Influenza Research and Surveillance (PGT) contract number HHSN272201400006C, the St. Jude Center of Excellence for Influenza Research and Response (PGT) contract number 75N93021C00016, 3U01AI144616-02S1 (PGT), and R01AI136514 (PGT). The Ellebedy laboratory was supported by NIAID grants U01AI141990 and U01AI150747, NIAID Centers of Excellence for Influenza Research and Surveillance contracts HHSN272201400006C and HHSN272201400008C, and by NIAID Collaborative Influenza Vaccine Innovation Centers contract 75N93019C00051. The contents of this publication are solely the responsibility of the authors and do not necessarily represent the official views of the NIAID or NIH.

## Author contributions

Conceptualization: PAM, AAM, MVP, PGT, AHE; Methodology: PAM, AAM, MP, JST, JP, JCC, MKK, SAT, JAO, RMP, PGT, AHE; Investigation: PAM, AAM, MVP, JST, WK, EK, JP, AJS, TL, AH, AMK, RCM, JCC, THON, LCR, ER, TS, WDM, SAT; Formal analysis: PAM, AAM, MVP, JCC; Visualization: PAM, AAM, MVP; Resources: SJTRC Study Team; Data curation: JCC, JW, SJTRC Study Team; Funding acquisition: PAM, PGT, AHE; Supervision: PAM, SAT, JAO, RMP, KK, JR, PGT, AHE; Writing – original draft: PAM, AAM, MVP; Writing – review & editing: All co-authors

## Methods

### Human cohort and HLA typing

Human subjects who elected to receive the BNT162b2 mRNA vaccine were recruited into this prospective observational study. Written informed consent was obtained from each subject. The study was approved by the Washington University in St. Louis Institutional Review Board (approval # 2020-12-081). Details of the cohort, including details regarding FNA sample collection, have been previously reported (Turner et al., 2021). Briefly, draining axillary lymph nodes ipsilateral to the deltoid vaccination site were located with ultrasound and sampled with multiple passes of 6 separate 25-gauge needles under real-time ultrasound guidance (Turner et al., 2020). Each needle was flushed with 3 mL of R10 (RPMI 1640 supplemented with 10% FBS and 100 U/mL penicillin-streptomycin) and the 3 separate 1 mL rinses of R10. Red blood cells were lysed with 1xACK (Sacha and Watkins, 2010) and then washed with P2 (1xPBS supplemented with 2% FBS and 2 mM EDTA). FNA samples were immediately stained for flow cytometry or cryopreserved in freezing media (10% dimethylsulfoxide and 90% FBS). Two subjects – 07 and 15 – received their BNT162b2 vaccine in the contralateral arm to the initial axillary lymph node FNA site. Subject 15 then had FNA performed on two lymph nodes, one ipsilateral and the other contralateral to the deltoid vaccination for all FNA samples completed starting on day 28. Matched blood samples from the same time-points were obtained by standard phlebotomy into EDTA anti-coagulated tubes and PBMC were prepared by density gradient centrifugation over Ficoll 1077 (GE). PBMC were treated with 1xACK for 5 minutes to lyse residual red blood cells before washing with R10 and immediate use in flow cytometry experiments or cryopreservation in freezing media.

For S167-180 tetramer validation and ICS experiments we used PBMC from SARS-CoV-2 convalescent and vaccinated donors obtained as a part of the St. Jude Tracking of Viral and Host Factors Associated with COVID-19 study (SJTRC, NCT04362995); a prospective, IRB-approved, longitudinal cohort study of St. Jude Children’s Research Hospital adult (≥18 years old) employees. Participants were screened for SARS-CoV-2 infection by PCR approximately weekly when on St. Jude campus. For this study, we utilized the convalescent blood draw for SARS-CoV-2 infected individuals (3-8 weeks post diagnosis) as well as post-vaccination blood draws for SARS-CoV-2 naive individuals. Blood samples were collected in 8 mL CPT tubes; and PBMC was isolated and frozen within 24 hours of collection. HLA typing of each participant was performed using the AllType NGS 11-Loci Amplification Kit (One Lambda; Lot 013) according to manufacturer’s instructions. Resulting libraries were sequenced on MiSeq lane at 150×150bp. HLA types were called using the TypeStream Visual Software from One Lambda.

### Cell sorting and flow cytometry

Fresh or frozen PBMC and/or FNA samples were washed and re-suspended in P2. For sorting of TFH populations from frozen FNA samples in Figures 1C and 5, cells were stained with CD4 Alexa Fluor 700 (SK3, BioLegend), CD19 PE (HIB19, BioLegend), CXCR5 PE-Dazzle 594 (J252D4, BioLegend), PD1 BB515 (EH12.1, BD Horizon), and Zombie Aqua (BioLegend) for a total of 30 minutes on ice. Cells were then washed twice with P2 and live, singlet, CD4^+^CD19^-^CXCR5^+^PD1^+^ cells were sorted on a FACSAria II into Trizol before being immediately frozen on dry ice.

In the bulk lymph node TFH versus spike-specific germinal center B cell experiment, FNA samples were stained in P2 for 30 minutes on ice with biotinylated and Alexa Fluor 647 conjugated recombinant soluble Spike proteins as well as PD-1 BB515 (EH12.1, BD Horizon). Cells were then washed twice with P2 and stained with IgG BV480 (goat polyclonal, Jackson ImmunoResearch), IgA FITC (M24A, Millipore), CD45 A532 (HI30, Thermo), CD38 BB700 (HIT2, BD Horizon), CD20 Pacific Blue (2H7, BioLegend, CD27 BV510 (O323, BioLegend), CD8 BV570 (RPA-T8, BioLegend), IgM BV605 (MHM-88, BioLegend), HLA-DR BV650 (L243, BioLegend), CD19 BV750 (HIB19, BioLegend), CXCR5 PE-Dazzle 594 (J252D4, BioLegend), IgD PE-Cy5 (IA6-2, BioLegend), CD14 PerCP (HCD14, BioLegend), CD71 PE-Cy7 (CY1G4, BioLegend), CD4 Spark685 (SK3, BioLegend), streptavidin APC-Fire750 (BioLegend), CD3 APC-Fire810 (SK7, BioLegend) and Zombie NIR (BioLegend) diluted in Brilliant Staining buffer (BD Horizon). Cells were then washed twice more with P2, fixed with the True Nuclear fixation kit (BioLegend) for 1 hour at room temperature, washed twice with True Nuclear Permeabilization/Wash buffer and then stained for 1 hour at room temperature with FoxP3 BV421 (206D, BioLegend), Ki-67 BV711 (Ki-67, BioLegend), Tbet BV785 (4B10, BioLegend), Bcl6 PE (K112-91, BD Pharmingen) and BLIMP1 Alexa Fluor 700 (646702, R&D Systems). Cells were then washed twice with True Nuclear Permeabilization/Wash buffer before acquisition on a Cytek Aurora spectral flow cytometer using SpectroFlo v2.2 software (Cytek) and analyzed using FlowJo software (v10.8.0, BD). Correlation in Figure 1B was performed in Prism (v9.1.0, GraphPad Software, LLC).

In tetramer staining experiments cells were stained in P2 for 10 minutes on ice with PE-labeled HLA-DPB1*04:01 S167-180 tetramer. Then, without washing away the tetramer, a master mix was added to the cells that included pre-titrated volumes of the following reagents: CD8 BV570 (RPA-T8, BioLegend) CD3 APC-Fire 810 (SK7, BioLegend) CD4 Alexa Fluor 700 (SK3, BioLegend) CD45RO BV650 (UCHL1, BioLegend) CCR7 BV785 (G043H7, BioLegend) CXCR5 PE-Dazzle 594 (J252D4, BioLegend) PD1 BB515 (EH12.1, BD Horizon) HLA-DR BV605 (L243, BioLegend) CD38 BB700 (HIT2, BD Horizon) ICOS BV421 (C398.4A, BioLegend) CD27 BV510 (O323, BioLegend) CD19 BV750 (HIB19, BioLegend) CD20 Pacific Blue (2H7, BioLegend) IgD PE-Cy7 (IA6-2, BioLegend) Zombie NIR (BioLegend) Spike protein conjugated to Alexa 647 and Spike protein conjugated to Alexa 488 and Brilliant Staining buffer (BD Horizon). Samples were then incubated on ice for an additional 30 minutes before they were washed twice with P2 and fixed in a final concentration of 1% paraformaldehyde for 15 minutes at room temperature. Samples were then run on a Cytek Aurora spectral flow cytometer using SpectroFlo v2.2 software (Cytek) and analyzed using FlowJo software (v10.8.0, BD). Tetramer responses over time in Figures 3 and 4 were graphed in Prism (v9.1.0, GraphPad Software, LLC).

### Jurkat cell line generation

For Jurkat cell line generation we selected a TCRα (TRAV35, CAGMNYGGSQGNLIF, TRAJ42) and two different TCRβ chains (TRBV4-1, CASSQGVGYTF, TRBJ1-2; TRBV6-3, CASSYRGAYGYTF, TRBJ1-2) from Bacher et al. Both TCRα and TCRβ chains were modified to use murine constant regions to facilitate surface expression (murine TRAC*01 and murine TRBC2*01). Two gBlock gene fragments were synthesized by Genscript to encode the modified TCRα chain, one of the modified TCRβ chains, and mCherry fluorescent protein, linked together by 2A sites. These sequences were cloned into the pLVX-EF1α-IRES-Puro lentiviral expression vector (Clontech). To generate the lentivirus we transfected 293T packaging cell line (ATCC CRL-3216) with the pLVX lentiviral vector containing TCR_4.1-mCherry or TCR_6.3-mCherry insert, psPAX2 packaging plasmid (Addgene plasmid #12260), and pMD2.G envelope plasmid (Addgene plasmid #12259). Viral supernatant was collected and concentrated using Lenti-X Concentrator 24- and 48-hours after the transfection (Clontech). Jurkat 76.7 cells (a gift from Wouter Scheper; variant of TCR-null Jurkat 76.7 cells that expresses human CD8 and an NFAT-GFP reporter) were transduced, then antibiotic selected for 1 week using 1 µg/mL puromycin in RPMI (Gibco) containing 10% FBS and 1% penicillin/streptomycin. Transduction of Jurkat cell line was confirmed by expression of mCherry, and surface TCR expression was confirmed via flow cytometry on BD Fortessa using FACSDiva software using antibodies against mouse TCRβ constant region (APC-Fire750-conjugated, Biolegend 109246), clone H57-597) and human CD3 (Brilliant Violet 421-conjugated, Biolegend 344834, clone SK7). Flow data was analysed in FlowJo 10.7.1.

### Jurkat peptide stimulation

Jurkat 76.7 cells expressing TCRs 4.1 and 6.3 (2.5×10^5^) were co-cultured with PBMCs from SARS-CoV-2 naive DPB1*:04:01-positive donor (6×10^5^) pulsed with 1 µM of CTFE**YVSQPFLMD**LE peptide, 1 µg/mL each of anti-human CD28 (BD Biosciences 555725) and CD49d (BD Biosciences 555501). An unstimulated (CD28, CD49d) and positive control (CD28, CD49d, 1X Cell Stimulation Cocktail, PMA/ionomycin; eBioscience 00-4970-93) were included in each assay. Cells were incubated for 18 hours (37 °C, 5% CO2). After the incubation cells were washed twice with FACS buffer (PBS, 2% FBS, 1 mM EDTA), resuspended in 50 µL of FACS buffer, and then blocked using 1 µL human Fc-block (BD Biosciences 564220). Cells were then stained with 1 µL Ghost Dye Violet 510 Viability Dye (Tonbo Biosciences 13-0870-T100) and a cocktail of fluorescent antibodies: 1 µL each of anti-human CD3 (Brilliant Violet 421-conjugated, Biolegend 344834, clone SK7), anti-human CD69 (PerCP-eFluor710-conjugated, eBioscience 46-0699-42, clone FN50), and anti-mouse TCRβ chain (APC/Fire750-conjugated, Biolegend 109246, clone H57-597). Cells were incubated for 20 minutes at room temperature and then washed with a FACS buffer. Cells were analyzed by flow cytometry on a custom-configured BD Fortessa using FACSDiva software (Becton Dickinson). Flow cytometry data were analyzed using FlowJo 10.7.1 software (TreeStar). Responsiveness to peptide stimulation was determined by measuring frequency of NFAT-GFP, CD69 and αβTCR expression.

### Peptide stimulation and intracellular cytokine staining of SJTRC samples

Donor PBMCs were thawed, suspended in RPMI 1640 supplemented with 10% human AB serum (Gemini Bio-Products, 100-512), 100 U/mL penicillin-streptomycin, 1% non-essential amino acids (Gibco, 11140-050), and 1 mM sodium pyruvate (Gibco, 11360-070), and plated at 2.5-4.0×10^5^ cells/well in a 96-well plate. PBMCs were stimulated with 5 µg/mL CTFE**YVSQPFLMD**LE peptide or left unstimulated and incubated at 37°C and 5% CO2. After 12 h, 1x PMA/ionomycin (eBioscience, 00-4970-93) was added to positive control wells and GolgiPlug (BD, 555029) was added at 1:1000 to all wells. Cells were incubated for an additional 6 h (for 18 h total), washed twice with FACS (PBS, 2% FBS, 1 mM EDTA), resuspended in 50 µL FACS containing 5 µL human Fc-block (Biolegend, 422302), and blocked for 15 min at RT. Cells were surface stained in an additional 50 uL FACS containing 1 µL Ghost Dye Violet 510 Viability Dye (Tonbo Biosciences 13-0870-T100) and a cocktail of fluorescent anti-human antibodies: CXCR5 SuperBright 436 (Thermo, 62-9185-42, clone MU5UBEE), CD45RA eFluor 450 (Thermo, 48-0458-42, clone HI100), CD19 BV510 (Biolegend, 302242, clone HIB19), CD8 BV570 (Biolegend, 301038, clone RPA-T8), CD3 BV750 (Biolegend, 344846, clone SK7), CD4 BB515 (BD, 565996, clone SK3), PD1 FITC (Biolegend, 329904, clone EH12.2H7), ICOS PerCP/Cy5.5 (Biolegend, 313518, clone C398.4A), CD69 PE/Cy7 (Biolegend, 310912, clone FN50), and γδ TCR AlexaFluor 647 (Biolegend, 331214, clone B1) for 30 min at 4°C. Cells were washed twice with FACS, fixed in Fix/Perm Solution (BD, 554715) for 20 min at 4°C, and washed twice in Wash/Perm buffer (BD, 554715). For detection of intracellular cytokines, cells were resuspended in 50 µL Perm/Wash buffer containing a cocktail of anti-human antibodies including IFNγ BV480 (BD, 566100, clone B27), TNFα BV605 (Biolegend, 502936, clone MAb11), IL17 BV785 (Biolegend, 512338, clone BL168), IL21 PE (Biolegend, 513004, clone 3A3-N2), and IL2 APC (Thermo, 17-7029-82, clone MQ1-17H12) and were incubated for 30 min at 4°C. Cells were washed twice in FACS and analyzed by flow cytometry on a Cytek Aurora spectral flow cytometer using SpectroFlo v2.2 software (Cytek) and analyzed using FlowJo v10.7.1 software (TreeStar). Responsiveness to peptide stimulation was determined by comparing the number of activated CD4^+^(CD69^+^) T cells positive for IL2, IFNγ, or TNFα production per 10^6^ PBMCs to matched unstimulated controls.

### Monomer generation

HLA-DP4 monomers with the S167-180 epitope were produced from purified HLA-DP4 containing the class II-associated invariant chain peptide (CLIP) (Niehrs et al., 2019) via HLA-DM catalysed peptide exchange as described previously for HLA-DR (Scally et al., 2013). Briefly, HLA-DP4 CLIP was expressed in Trichoplusia Ni (Hi5) insect cells *via* a pFastBac-Dual construct encoding HLA-DP4 α- and β- chains with C-terminal fos/jun zipper domain. The HLA-DP4 β-chain further contained an N-terminal factor Xa cleavable CLIP sequence, and a C-terminal biotinylation signal and His7 tag (Niehrs et al., 2019). Following expression for 3 days at 27 °C, cell supernatant was concentrated and buffer exchanged in a Tangential Flow Filtration system into 500 mM NaCl, 10 mM Tris-HCl pH8 and subsequently purified *via* immobilised metal affinity chromatography and Superdex S200 gel permeation chromatography (GPC) in 150 mM NaCl, 10 mM Tris-HCl pH8. The linked CLIP peptide was cleaved with factor Xa for 6 h at 21°C prior to peptide exchange, and factor Xa cleaved HLA-DP4 was subsequently incubated in the presence of a 10-fold molar excess of peptide and a 1/5 molar ratio of HLA-DM for 16h at 37°C in 100 mM sodium citrate pH 5.4. HLA-DP4 loaded with S167-180 peptide was buffer exchanged into 50mM NaCl, 20 mM Tris-HCl pH8, purified via Hi-Trap Q ion exchange chromatography and biotinylated using BirA biotin ligase. Following a final Superdex S200 GPC step in PBS, biotinylated HLA-DP4-S167-180 monomer was concentrated to approx. 1mg/ml and stored at -80 °C.

### Tetramer generation and staining of Jurkat cells

Biotinylated DPB1*04:01-monomers loaded with TFE**YVSQPFLMD**LE peptide (S167-180) were tetramerised using PE-Streptavidin (Biolegend, 405204). 1.1 volume PE-conjugated streptavidin was added to 1 volume of DPB1*04:01-monomer (1 mg/mL). The volume of PE-streptavidin (0.2 mg/ml) was divided in 4 parts and added in 4 consecutive steps with 10 minutes incubation between. After adding all needed amounts of PE-streptavidin the mixture was incubated for at least 1 hour on ice prior to staining. Jurkat 76.7 cells expressing TCR4.1, TCR6.3, Jurkat 76.7 cell line expressing irrelevant TCR (specific to NQKLIANQF epitope from the spike protein of SARS-CoV-2, REF), and SARS-CoV-2 naive DPB1*04:01 positive donors’ PBMCs were stained with 1 µL Ghost Dye Violet 510 Viability Dye (Tonbo Biosciences 13- 0870-T100) and 1 µL of DPB1*04:01-S167-180-tetramer. Cells were analyzed by flow cytometry on a custom-configured BD Fortessa using FACSDiva software (Becton Dickinson). Flow cytometry data were analyzed using FlowJo 10.7.1 software (TreeStar). The quality of the S167-180 HLA class II tetramer was judged by staining of the relevant T cell line and low background in irrelevant Jurkats and naive PBMCs.

### Tetramer staining of SJTRC samples and scTCR sequencing

Donor PBMCs were thawed and resuspended in 100 µL FACS buffer (PBS, 0.5% BSA, 2 mM EDTA). Cells were stained with 5 µL Fc-block (Human TruStain FcX, Biolegend 422302) and 1.5 µL of S167-180 HLA class II PE-conjugated tetramer for 30 minutes on ice. After the incubation a cocktail of fluorescently-labeled surface antibodies (2 µL of each: Ghost Dye Violet 510 Viability Dye, Tonbo Biosciences 13-0870-T100; anti-human CD3 PerCP Cy5.5-conjugated (Biolegend 317336, clone OKT3), anti-human CD4 BV711-conjugated (Biolegend, 317440, clone OKT4), anti-human CD45RA BV421-conjugated (Biolegend, 304130, clone HI100), and anti-human CCR7 FITC-conjugated (Biolegend, 353216, clone G043H7) was added. Samples were incubated for an additional 20 minutes on ice. Single, Live, CD3-positive, CD4-positive, tetramer-positive cells were sorted on the Sony SY3200 into 384-well plates with premixed SuperScript VILO cDNA Synthesis mix (Invitrogen, 11754250) for subsequent scTCR sequencing. scTCR library preparation and sequencing was performed as previously described (Wang et al., 2012). In brief, cDNA underwent two rounds of nested multiplex PCR amplification with a forward primer mix specific for V-segments and reverse primers for C-segments of TCRalpha and TCRbeta and sequenced on Illumina MiSeq platform (2×150 read length). Resulting TCR sequences can be found in **Supplementary Table 1**.

### Bulk repertoire generation

TCRalpha and TCRbeta bulk repertoires were generated with the 5’RACE protocol described in (Egorov et al., 2015). RNA was isolated using Trizol reagent (Invitrogen) according to the manufacturer’s protocol. All RNA was used for cDNA synthesis with SmartScribe kit (Takara), using template switch oligonucleotide and primers specific for TCRalpha and TCRbeta constant segments. cDNA was amplified in two rounds of PCR using Q5 high-fidelity polymerase (NEB). Adapters necessary for sequencing on the Illumina platform were introduced with the KAPA HyperPrep kit (Roche). Libraries were sequenced on Illumina MiSeq platform (2×150).

### TCR repertoire analysis

Bulk TCR repertoire data was demultiplexed and assembled into the UMI consensuses with *migec* (v. 1.2.7; with collision filter and force-overseq parameters set to 1) (Shugay et al., 2014). V and J-segment alignment, CDR3 identification and assembly of reads into clonotypes were performed with MiXCR (v. 3.0.3) with default parameters (Bolotin et al., 2015). Resulting processed repertoire datasets and reference to raw TCR repertoire sequencing data are available at GEO database (acc. GSE183393). Analysis of bulk repertoire data was performed using R language for statistical computing, with merging and subsetting of data performed using *data.table* package. *stringdist* and *igraph* R packages were used to build TCR similarity network, *gephi* software was used for TCR similarity networks layout and visualisation and *ggplot2* library for other visualisations.

### Public TCR repertoire datasets

TCRbeta dataset for MIRA class II peptide stimulation (ImmuneCODE MIRA release 002.1) was accessed via ImmuneACCESS database (Nolan et al., 2020). Processed single cell paired chain TCR datasets from ARTE assays after 6 and 24 hour stimulation with SARS-CoV-2 peptides were used as supplied by authors in original publications: Table S3 from (Bacher et al., 2020), Table S4A from (Meckiff et al., 2020).

---

**Supplementary Figure 1.**
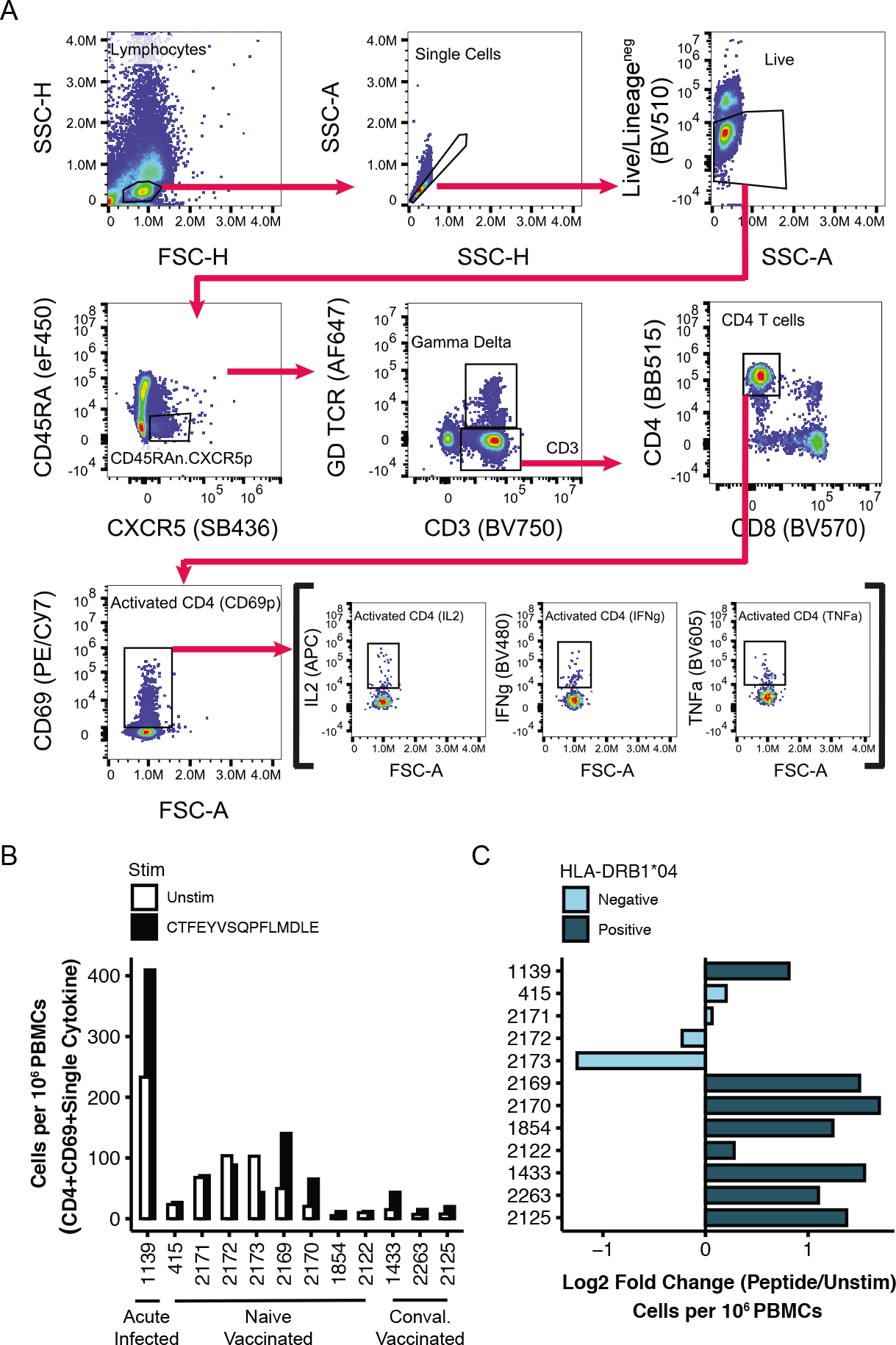
Intracellular cytokine staining of PBMCs stimulated with S166-180 peptide. **(A)** Gating strategy employed to resolve CD4^+^/CD69^+^ T cells producing IL2, TNFα, or IFNγ. Activated CD4^+^ T cells were defined as Live/B cell lineage (CD19^+^)^neg^/TFH lineage (CD45RA^-^/CXCR5^+^)^neg^/γδ TCR^neg^/ CD3^+^/CD4^+^/CD69^+^ and Boolean gated on IL2^+^, TNFα^+^, or IFNγ^+^ single-positive lymphocytes. **(B)** The number of CD4^+^/CD69^+^ T cells producing IL2, TNFα, or IFNγ per 10^6^ PBMCs following CTFEYVSQPFLMDLE peptide (black) or media (white) stimulation. **(C)** The log2-fold change of peptide-stimulated over unstimulated CD4^+^/CD69^+^ T cells producing IL2, TNFα, or IFNγ per 10^6^ PBMCs from DPB1*04:01/02- positive and -negative participants.

**Supplementary Figure 2.**
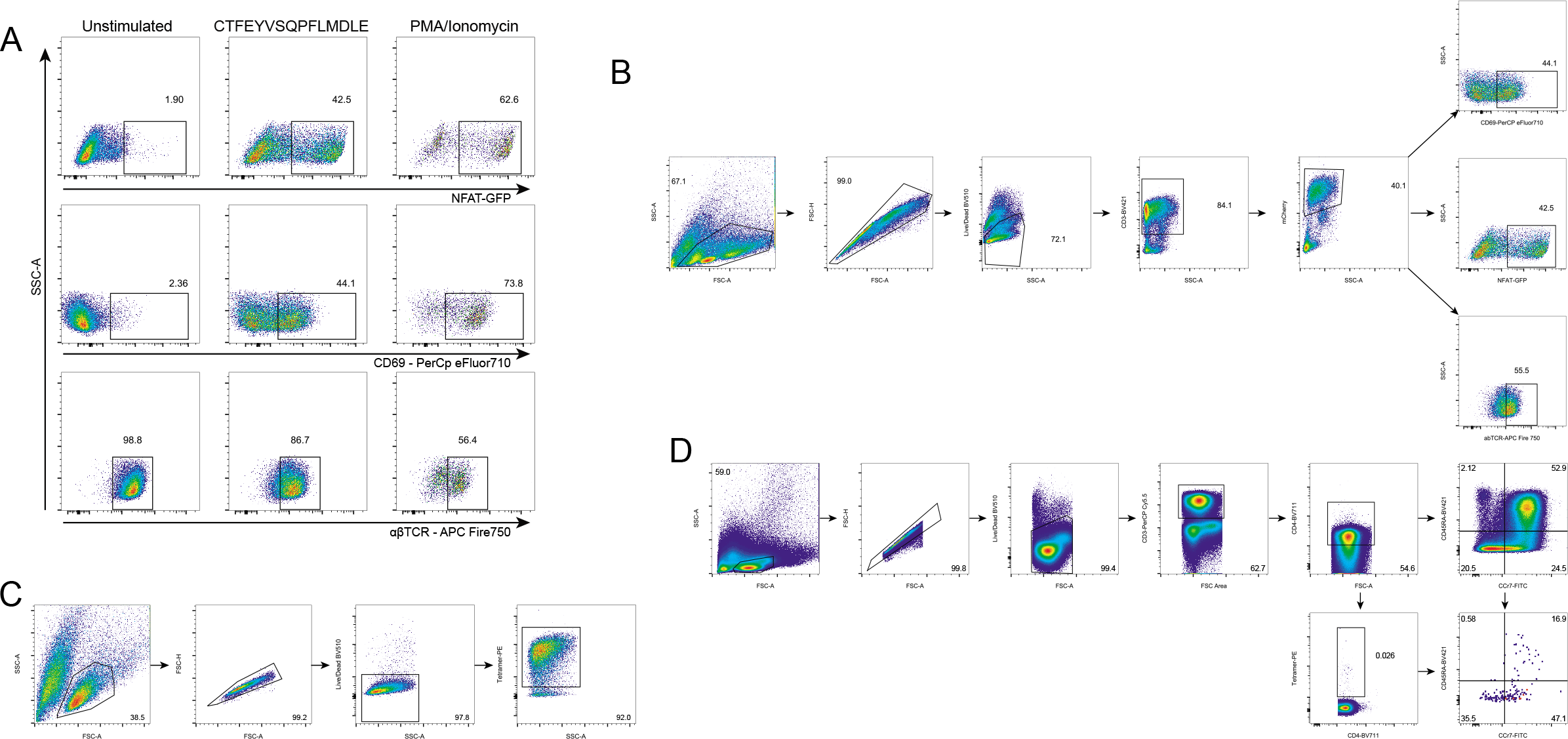
S167-180 epitope discovery and validation. **(A)** Jurkat cell line expressing the predicted TCR after stimulation with the predicted epitope. Left column: negative control; middle column: TCR6.3 cell line co-cultured with PBMCs from healthy DPB1*04:01-positive donor pulsed with CTFEYVSQPFLMDLE peptide (S166-180); right column: positive control. Top row: NFAT-GFP reporter expression. Middle row: CD69 surface expression. Bottom row: Downregulation of the TCR on cell surface. **(B)** Gating strategy for **(A)** and Figure 2D. **(C)** Gating strategy for Figure 2E. **(D)** Gating strategy for Figure 2F.

